# From temporal network data to the dynamics of social relationships

**DOI:** 10.1101/2021.03.22.436267

**Authors:** Valeria Gelardi, Alain Barrat, Nicolas Claidiere

## Abstract

Networks are well-established representations of social systems, and temporal networks are widely used to study their dynamics. Temporal network data often consist in a succession of static networks over consecutive time windows whose length, however, is arbitrary, not necessarily corresponding to any intrinsic timescale of the system. Moreover, the resulting view of social network evolution is unsatisfactory: short time windows contain little information, whereas aggregating over large time windows blurs the dynamics. Going from a temporal network to a meaningful evolving representation of a social network therefore remains a challenge. Here we introduce a framework to that purpose: transforming temporal network data into an evolving weighted network where the weights of the links between individuals are updated at every interaction. Most importantly, this transformation takes into account the interdependence of social relationships due to the finite attention capacities of individuals: each interaction between two individuals not only reinforces their mutual relationship but also weakens their relationships with others. We study a concrete example of such a transformation and apply it to several data sets of social interactions. Using temporal contact data collected in schools, we show how our framework highlights specificities in their structure and temporal organization. We then introduce a synthetic perturbation into a data set of interactions in a group of baboons to show that it is possible to detect a perturbation in a social group on a wide range of timescales and parameters. Our framework brings new perspectives to the analysis of temporal social networks.

## Introduction

Social relationships are created and maintained through interactions between individuals which last and are repeated over a variety of timescales. Social networks provide convenient representations for the resulting human and non-human animal social structures, where individuals are the nodes of the networks and links (ties) are summaries of their social interactions [Granovetter, 1973, Hinde, 1976, Wasserman and Faust, 1994, Brent et al., 2011]. Since the early definition of the sociogram [Moreno, 1934], these networks have typically been constructed by aggregating dyadic interactions occurring over a certain period of time to define the links between individuals. Research on the resulting static networks has led to numerous insights into human and non-human societies, with the development and empirical verification of various social theories such as the social balance hypothesis [Heider, 1946, Szell et al., 2010, Gelardi et al., 2019], the “strength of weak ties” theory [Granovetter, 1977, Karsai et al., 2014] or Dunbar’s theory on a cognitive limit to the possible number of simultaneous relationships [Dunbar, 1998, Gonçalves et al., 2011].

By definition however, such static networks do not capture the dynamics of social relationships within the aggregation period. As noted by Granovetter in 1973, further development of social network analysis requires “*a move away from static analyses that observe a system at one point in time and to pursue instead systematic accounts of how such systems develop and change*” [Granovetter, 1973]. Important advances in this respect have been made possible by the recent availability of temporally resolved data on interactions between individuals, from various types of communication [Eckmann et al., 2004, Kossinets and Watts, 2006, Onnela et al., 2007, Karsai et al., 2011, Miritello et al., 2013b] to face-to-face interactions [Cattuto et al., 2010, Salathé et al., 2010, Stopczynski et al., 2014, Toth et al., 2015]. These data fueled the development of the framework of temporal networks [Holme and Saramäki, 2012, Holme, 2015], which replaces static ties by information on the actual series of interactions on each tie.

Temporal data and temporal networks have allowed researchers to further the study of social networks in various ways. For instance, aggregating temporal information over successive time windows, has made it possible to follow the evolution of ties over larger timescales [Saramäki et al., 2014, Fournet and Barrat, 2014, Gelardi et al., 2019, Aledavood et al., 2015], revealing circadian interaction patterns [Aledavood et al., 2015], for example, or the stability of how individuals divide interaction time among their relationships, even in different periods of their lives and with different groups of friends [Saramäki et al., 2014]. The use of digital and phone communication data has yielded further insights into social theories, such as the various strategies individuals use to manage their social circle when faced with limited communication resources [Miritello et al., 2013a, Miritello et al., 2013b]. Taking into account the temporal features of each tie during a certain time window can also shed light on their strength and persistence [Navarro et al., 2017, Ureña-Carrion et al., 2020]. Finally, researchers have identified temporal structures with no static equivalent [Kovanen et al., 2011, Kobayashi et al., 2019, Galimberti et al., 2018] that can reveal interesting patterns of relevance to the analysis of social phenomena or dynamic processes in a social group [Kovanen et al., 2013, Ciaperoni et al., 2020].

Despite this wealth of studies and results, moving from a stream of interactions within a group of individuals, represented by a temporal network, to a meaningful representation of the evolution of their social relationships remains a challenge. Indeed, the temporal network seen at any specific time *t* contains by definition only the interactions taking place at t and a number of properties of the networks obtained by temporal aggregation on successive windows depend on the window length and placement [Sulo et al., 2010, Krings et al., 2012, Psorakis et al., 2012, Kivelä and Porter, 2015]. Aggregating over increasing time window lengths also averages out relevant temporal information and no single natural time scale for aggregation can be defined, as relevant dynamics occur on multiple timescales [Holme, 2013, Saramäki and Moro, 2015, Darst et al., 2016, Masuda and Holme, 2019].

Here, we address this issue by putting forward a new systematic way to transform the stream of interactions between individuals (the temporal network data) into a continuously evolving representation of the social structure, i.e., a network with time-varying weights. In other words, the evolving weight *w*_*ij*_(*t*) of the tie between nodes *i* and *j* should give information on the status of their relationship at *t*. To date, few such dynamic network models have been proposed [Ahmad et al., 2018, Zuo and Porter, 2019, Jin et al., 2001, Palla et al., 2007], mainly based on the idea that the weight of a tie between two individuals strengthens when they interact, and that in the absence of interaction, the tie’s weight decays exponentially with time (the timescale of the decay is the model’s parameter). However, these rules of evolution assume that the links between distinct pairs of individuals are independent, while the interdependence of social relationships is often well justified, especially for primates. Compared to most other animals, humans and other primates form complex social groups characterized by long-term relationships that are both structured and flexible [Dunbar and Shultz, 2007,Mitani, 2009, Silk et al., 2010]. The creation and maintenance of these relationships require specific cognitive skills [Cheney et al., 1986], for instance in helping others [Burkart et al., 2014] or understanding others’ intentions [Tomasello et al., 2005], and there is now strong evidence that the evolution of brain sizes in primates has been driven, at least in part, by the corresponding demands of social life [Dunbar and Shultz, 2007, Dunbar and Shultz, 2017, Lewis et al., 2011, Kwak et al., 2018, Noonan et al., 2018, Taebi et al., 2020, Meguerditchian et al., 2021]. Thus, in primates, investing in a social relationship is a costly strategic decision, controled by evolved cognitive mechanisms. The quality of an individual’s social relationships depends on the time invested in them [Dunbar, 2020, Dunbar et al., 2009] and has important life consequences. For instance, finite communication capacities can imply that the activation of a new social tie occurs at the expense of a previously existing one [Miritello et al., 2013a]. It is therefore crucial to take into account the finite capacities of each individual in establishing and maintaining social ties in order to represent the evolution of the weight of these ties. In particular, the occurrence of a social interaction between two individuals not only reinforces their mutual relationship, but it also weakens the relationships they have with others: the time and energy spent to maintain the tie with an individual is taken from a finite interaction capacity and thus is time that is not spent with others.

The framework that we put forward here uses this type of interdependence of social relationships to transform a stream of interactions into an evolving weighted network: with each interaction between two individuals, the weight of their tie increases, while the weight of the ties they have with other individuals decreases. In contrast to other recent temporal network representations [Ahmad et al., 2018, Zuo and Porter, 2019], time itself is not explicit, and the weight of a tie remains unchanged if the corresponding individuals do not interact with anyone. Our framework is therefore linked to the Elo rating method [Elo, 1978] used to rank chess players and analyze animal hierarchies: the dynamics of the system are determined by the pace of interactions between individuals, not by the absolute time between events.

In the following, we define a parsimonious model for the evolution of social ties based on these concepts, with two parameters quantifying respectively the increase in the weight of a tie *i* − *j* when an interaction occurs between *i* and *j*, and its decrease when another interaction involving either *i* or *j* (but not both) takes place. We then show the relevance of the model by applying it to several data sets describing interactions in groups of human and non-human primates and by using it to automatically detect naturally occurring changes in the groups’ dynamics and artificially generated perturbations in the data.

## Results

### Framework

The framework and concepts highlighted above can be translated in various ways into modeling rules to transform a stream of dyadic interactions into evolving weights on each tie of an evolving network *G*(*t*). The nodes of the network represent the individuals and the weight *w*_*ij*_(*t*) of tie *i* − *j* represents the strength of their social relationship at time t. More specifically, we here use a model in which *G(t)* is directed, i.e., *w*_*ij*_(*t*) represents the strength of the relationship *seen from i*, which is not necessarily equal to the strength seen from *j*, *w*_*ji*_(*t*). This reflects the fact that a relationship does not necessarily have the same importance for both individuals involved.

The model depends on two parameters, *α* and *β*, and evolves according to the following rules:

- We start from an empty network with uniform weights initialized to zero, i.e., *w*_*ij*_(0) = 0 ∀*i*, *j*;
- For each interaction between nodes *i* and *j* at time *t*, the weights of the ties in which *i* and *j* are involved are updated according to

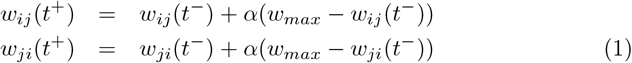

and

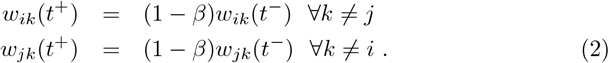

Here, *t*^−^ and *t*^+^ stand respectively for the times immediately before and after the interaction. The parameter 0 < *α* < 1 quantifies how much a tie strength is reinforced by each interaction, while 0 < *β* < 1 accounts for the weakening of the strength of the ties with other individuals. *w*_*max*_ > 0 represents the maximum possible value of the weights, which we set to *w*_*max*_ = 1 without loss of generality. These rules ensure that the weights all remain bounded between 0 and *w*_*max*_. They also mean that if a tie’s weight is zero, it remains so unless there is an interaction involving that tie, and that individuals who interact often see the weight of their tie increase towards *w*_*max*_.

It is important to stress once again that while instantaneous interactions may be undirected, i.e., there are no source nor target individuals (e.g. in face-to-face interaction data), the evolution rules (1)-(2) naturally result in a directed network. For instance in an interaction between *i* and *j*, the weight *w*_*ik*_ between *i* and an individual *k* ≠ *j* decreases because i devotes time to *j* but not to *k*, while the weight *w*_*ki*_ does not change.

The evolution rules could easily be modified in the case of directed interactions, such as in an exchange of text messages or on online social media: for instance, if *i* sends a message to *j*, the weights *w*_*ij*_ and *w*_*ik*_ could be affected more strongly than the weights *w*_*ji*_ and *w*_*jk*_. However, this would require the introduction of additional parameters.

Finally, we note that the evolution rules can be applied to temporal network data expressed either in continuous time (i.e., an interaction between two individuals can occur at any time) or in discrete time (when the data itself has a finite temporal resolution).

### Application to empirical data

Let us first consider the application of the framework described above to empirical data describing interactions in close proximity (as collected by wearable devices) in two schools, namely a French elementary school [Stehlé et al., 2011b] and a US middle school in Utah [Toth et al., 2015,Leecaster et al., 2016], with a temporal resolution of approximately 20 seconds in both cases (see Materials and Methods for more details on the data sets). Although both cases involve school contexts, the classes were organized very differently, as described in [Stehlé et al., 2011b, Leecaster et al., 2016]: the elementary school students remained in the same classroom for their different classes, while the middle school students changed classrooms between classes.

In each case, we transformed the temporal network data into a network of ties *G*(*t*) between individuals, with the weights evolving according to the rules (1)-(2). For simplicity, we used *α* = *β* and considered various values of *α*. We then stored the network *G*(*t*) and the tie weights every Δ time steps (i.e., we store *G*(*n*Δ) for *n* = 0, 1, 2,) and computed the similarity between each pair of the stored networks *G*(*n*Δ) and *G*(*n*′Δ) (see Materials and Methods). We thus obtained a matrix of similarity values [Masuda and Holme, 2019, Gelardi et al., 2019] for each value of *α*, shown in Figure 1 for *α* = 0.1 (see Figure S1 of the Supplementary Material for other values of *α*). These matrices clearly highlight that the two contexts correspond to different schedules and organizations of interactions. Moreover, in each case they reflect the temporal organization and reveal the various periods of importance in the school schedules.

**Figure 1.**
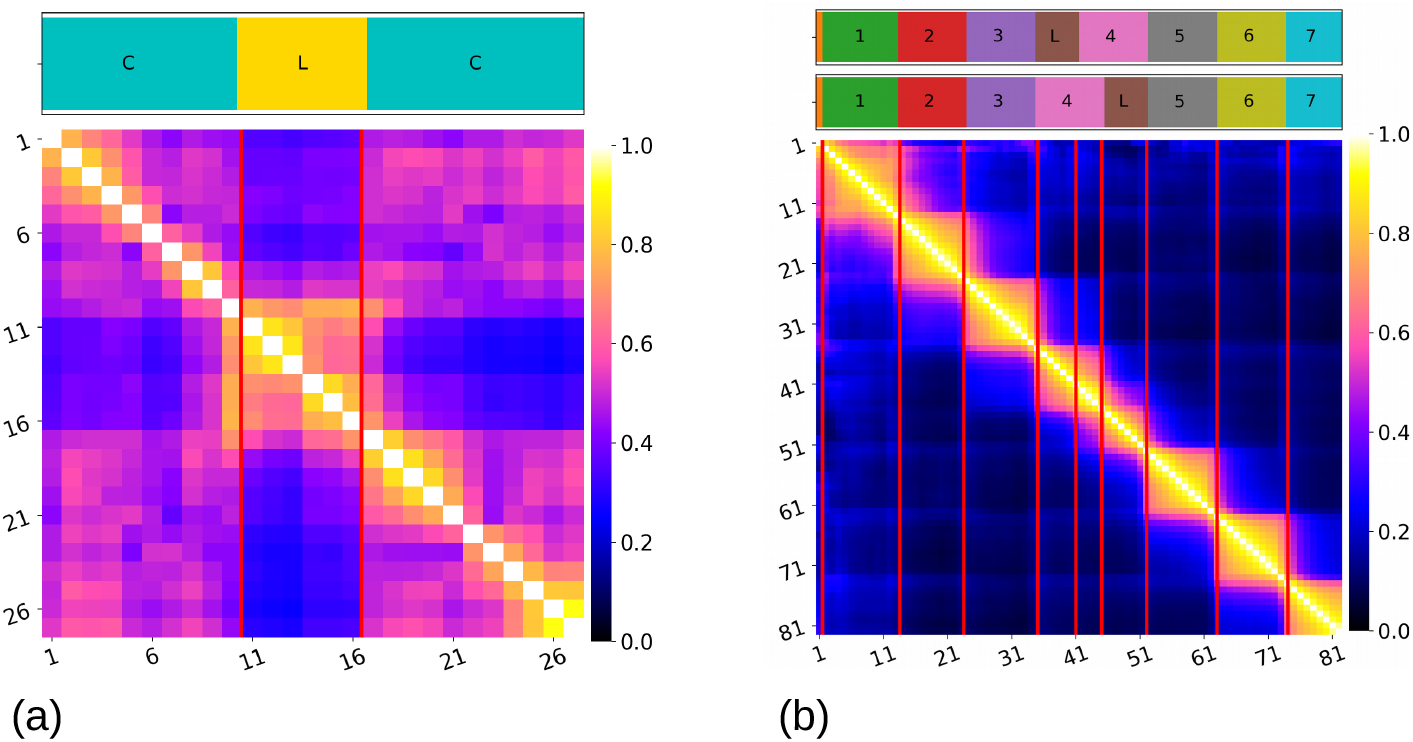
Similarity matrices and school schedules between the evolving networks built from the first day of data collected in the French elementary school (a) and the US middle school (b). Here we use *α* = *β* = 0.1, and the evolving networks are observed every Δ = 20 minutes for the French school and every Δ = 5 minutes for the US school. The horizontal bars at the top of the figures give information about the schedule of a school day. The different colors in the bar correspond to the different class times (indicated by the letter C in (a) and with different numbers in (b)) and lunchtimes (indicated by the letter L), the length of each colored interval representing the duration of the corresponding period. In (b) there are two bars because the students were split into two groups for their lunchtime and fourth class period and therefore have slightly different schedules. The vertical red lines indicate the beginning and the end of lunchtime in (a) and the starting times of the different classes and lunch periods in (b).

In the case of the French elementary school, the similarity between the networks at various times during each half-day is relatively high. The networks obtained during the lunch period (highlighted in the figure) are dissimilar to the networks obtained during class times, and a transition between two periods can be observed during lunch, in agreement with the description in [Stehlé et al., 2011b], which notes that students ate lunch in two successive groups. The networks obtained in the afternoon are similar to the ones from the morning, which is consistent with the fact that students returned to the same classrooms with the same seating arrangements.

The similarity matrix obtained for the networks representing the US middle school is strikingly different: here a strong similarity can be observed between networks during successive periods of time (yellow blocks along the diagonal, indicating high similarity and hence stable networks), with a very low similarity between networks observed in different periods. By examining the class schedules detailed in [Leecaster et al., 2016], we observe that each period of network stability indeed corresponds to a class or lunch period (see the colored bar at the top of Figure 1b). Note that the stability of the network in these periods is not due to a lack of interactions, as Figure S2 in the Supplementary Material makes clear. Moreover, the low similarity between different class periods can be understood from the fact that students switched classrooms between classes, in contrast with the elementary school students.

Examining the similarity matrices obtained from the weighted evolving networks thus provides important insights into the evolution of the systems under scrutiny and makes it possible to distinguish the occurrence of moments of stability and change in the structure of the network. While we used *α* = 0.1 in Figure 1, we considered other values in Figure S1 in the Supplementary Material, revealing that the distinction between the various periods is blurred for small values of *α* but becomes more and more apparent as *α* increases. In this figure, we show the similarity matrices corresponding to the full two days of data. For *α* = 0.1 the data highlight how the two lunch periods at the French elementary school are different from each other, while the class periods during the two days are similar. For the US middle school, we also observe a similarity between class periods during the two different days, reflecting the similarity of class schedules during these two days and indicating that the seating arrangements in each classroom were probably similar on different days. Finally, Figure S3 in the Supplementary Material displays the similarity matrices between temporal networks aggregated over time windows of different lengths, similarly to the procedure in [Masuda and Holme, 2019], where the distinction between lunch and class periods in the elementary school (see also [Masuda and Holme, 2019]) and between the middle school class periods is also observable.

### Detection of a perturbation

To go beyond a mere visual inspection of the similarity matrices, we considered a more systematic analysis of the capacity of a temporal network representation, obtained either by temporal aggregation or through our framework, to detect perturbations in a social group’s interaction patterns.

To this aim, we first introduced a synthetic perturbation of controled intensity and duration in the temporal network data, for instance by switching the identity of some nodes for a certain duration. We then followed the steps outlined in Fig. 2. First, we used our framework to transform the perturbed temporal network into an evolving weighted graph according to the evolution rules (1)-(2). This weighted graph was observed every *p* time steps (if the real time duration of one time step is *δ*, this means that we observed the graph every Δ = *pδ*). As a baseline, we also aggregated the temporal network data on successive time windows of duration Δ (Fig. 2a). We then followed Masuda et al.’s procedure for detecting states in a temporal network [Masuda and Holme, 2019]. Namely, we computed the cosine similarity matrix between graphs observed at different times (Fig. 2b) and transformed it into a distance matrix. We then applied a hierarchical clustering algorithm (see Material and Methods) in order to detect discrete states of the network. As the ground truth perturbation is known, we added a validation step to the procedure to compare the states obtained by the clustering algorithm to the perturbation timeframe. In this step we quantified the detection performance through two indicators (Fig. 2d), namely the Jaccard index between the sets of timestamps of the actual perturbation and the timestamps of the perturbed state detected, and the delay between the start time of the actual perturbation and the corresponding value obtained through the clustering algorithm (see Material and Methods).

**Figure 2.**
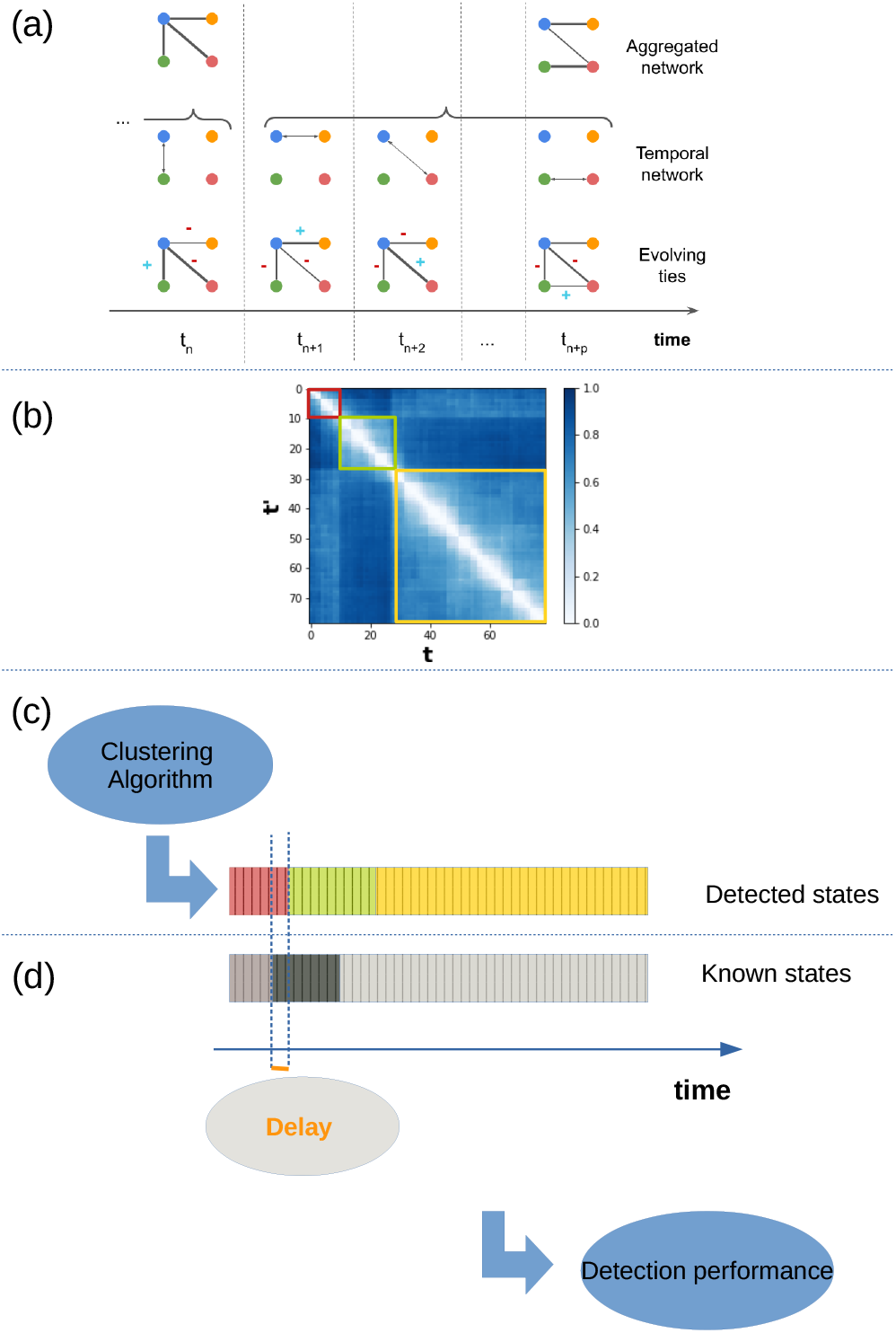
Workflow used to detect discrete states and change points in temporal networks (see also [Masuda and Holme, 2019]) and to estimate the performance of the detection. (a) Creation of a sequence of networks, either by temporal aggregation over successive time windows of *p* time steps, or by transforming the data into an evolving network observed every *p* time steps. (b) Computation of the similarity between all pairs of networks using the global cosine similarity (see Methods). (c) Classification of the networks into discrete states using a hierarchical clustering algorithm on the distance matrix (the distance between two graphs being simply defined as 1 minus their similarity). (d) Estimation of the performance of the classification obtained by the clustering by comparison with the ground truth perturbation time window using the Jaccard index between the actual and detected time frame of the perturbation and the shifts between actual and detected start times of the perturbation.

To illustrate the procedure, we considered proximity data from a group of 13 Guinea baboons (*Papio papio*), collected from June to November 2019 using wearable sensors with a temporal resolution of 20 seconds (see Material and Methods). We introduced a small perturbation in the data, namely the exchange of two individual’s identities in the data during a certain period. In Figure 3 we use a perturbation duration of 2 hours and show the resulting similarity matrices between the weighted evolving networks obtained for three values of *α* = *β* and observed every 30 minutes. We also measure and show the detection performance as a function of *α*. Strikingly, even such a small and short perturbation is well detected over a wide range of *α* values, excepting the smallest and largest. The perturbation is not detected for small *α* values, as the resulting network dynamics is too slow: Fig. 3(a) shows that the network remains very similar to itself during the whole explored time range. However, we observe a sharp increase in detection performance as soon as the resulting dynamics are fast enough. At very large *α* values, the detection becomes impossible again because each single interaction induces large changes in the weights, leading to rapidly changing dynamics with no stable period for the weighted evolving network. Overall, the perturbation is well detected over a wide range of *α* values. Notably, the perturbation is instead not detected when using temporal aggregation over successive windows of 30 minutes.

**Figure 3.**
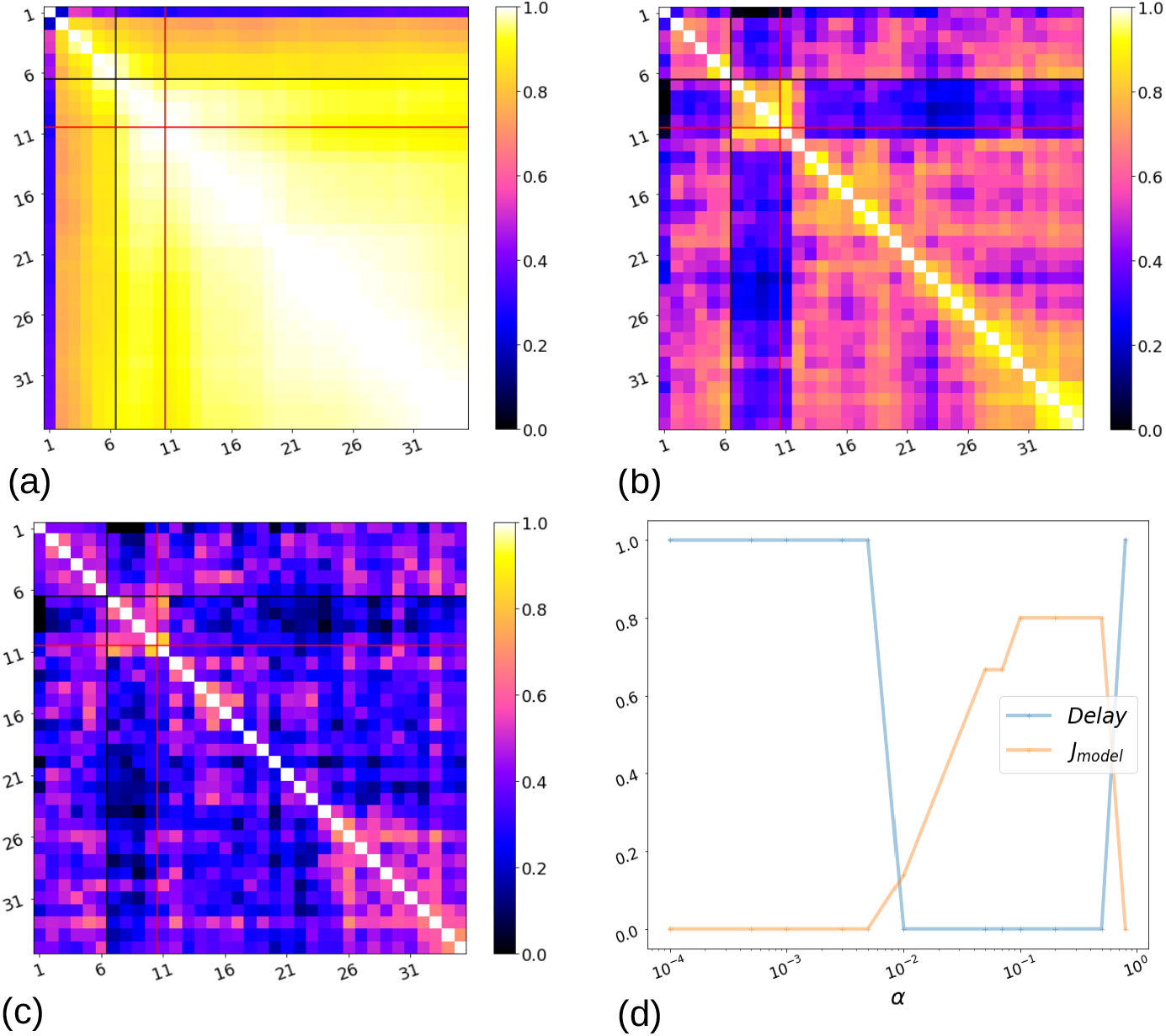
Detection of a simulated perturbation in a temporal network data set. Here we consider one day of proximity data collected from a group of 13 baboons (see Material and Methods). The data, with a temporal resolution of 20 seconds, are artificially perturbed by exchanging the identity of two nodes for 2 hours. The resulting perturbed temporal network is transformed into a weighted evolving network as described in the text, and this network is observed here every 30 minutes. Panels (a), (b), (c) represent the resulting cosine similarity matrices for values of *α* = *β* = 0.001, 0.1, 0.5, respectively. The black and red lines correspond to the (known) start and end times of the perturbation. Panel (d) shows the performance detection of network states (see Fig. 2), computed from the hierarchical clustering analysis applied to the distance matrices, with the number of clusters fixed to *C* = 3. The blue line represents the relative delay in the detection of the perturbation, i.e. the difference between the known beginning of the perturbation (black line) and the detection of a new network state, divided by the total length of the perturbation. The orange line indicates the Jaccard index between the known perturbation timestamps and the perturbation detected by the clustering algorithm. The detection performance relative to the aggregated network is not presented because no cluster detected by the algorithm could correspond to the simulated perturbation.

We also considered other time scales of perturbation and observation of the evolving networks (or aggregation of the temporal network): in Supplementary Figures S5 and S6, we illustrate these results for the same data set and for two different timescales. In Figure S5, we studied the evolution of the system over 20 days, observing the evolving network on a daily basis. We simulated a perturbation by switching the same two individuals as for Figure3 for 3 days. At such a timescale our framework results in a perfect or almost perfect detection of the perturbation for a wide range of values of the parameter *α* (i.e., values of the Jaccard index close or equal to 1), while the perturbation was not detected when using daily aggregated networks (Jaccard index equal to 0). In Figure S6, we used the entire period of data collection (from June to November 2019), and observed the evolving weighted network on a weekly basis. We perturbed the network, switching the same individuals as in the previous cases, for a period of 15 days, affecting weeks 6 to 8 (the perturbation started exactly in the middle of week 6; the networks were affected for 3 successive weeks). In this case, using a weekly aggregated network also made it possible to detect the perturbation, but the detection performance of our framework was higher for values of the parameter *α* larger than 0.001. Overall, even when the simple temporally aggregated networks are able to detect the perturbation, there is always a range of values of the model’s parameter *α* such that the evolving network representation provides a better detection performance.

We further investigated whether using different values for the parameters *α* and *β* could lead to an improvement in the detection performance. We show the results in Figure 4 for the same data and perturbation as for Figure 3 (see also Supplementary Figure S7). We found that the detection performance worsened for *β* < *α*, while it increased for *β* > *α*. The faster decay of ties induced by the larger value of *β* was indeed then able to compensate for dynamics which were too slow when obtained with small values of *α*: the change in interactions due to the perturbation were translated very quickly into the evolving weights. For instance, if a node *i* was repeatedly interacting with a node *j* before the perturbation, but interacts more with another one, *k*, during the perturbation, *w*_*ij*_ decreases quickly as soon as the perturbation starts, and this can be easily detected even if *w*_*ik*_ only increases slowly.

**Figure 4.**
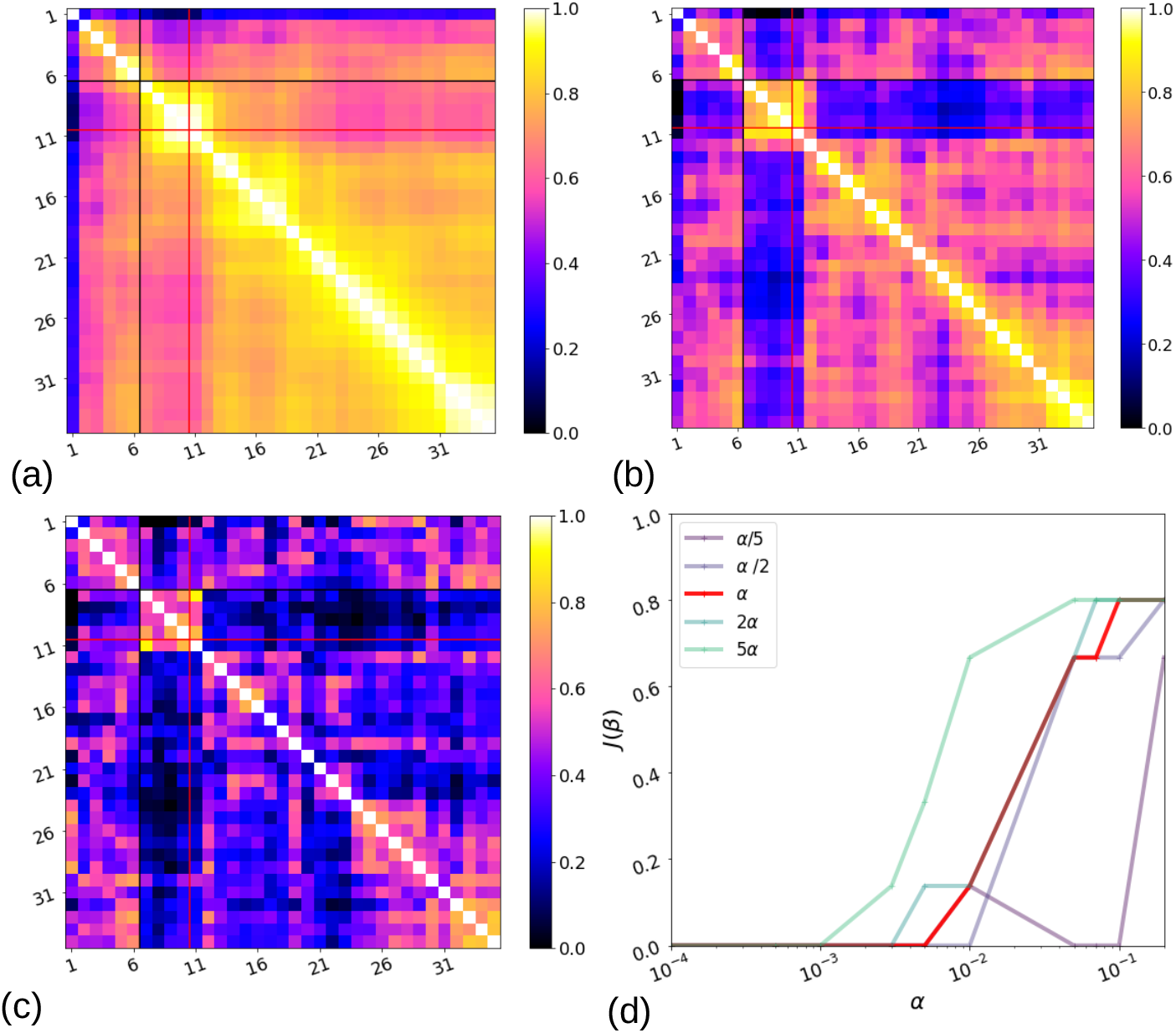
Performance of the detection of simulated changes when varying. *β*. Panels (a), (b), (c) represent the cosine similarity matrices for *α* = 0.1 and values of *β* = *α*/5, *α*, 5α, respectively, using the same simulated perturbation as in Fig. 3. Panel (d) shows the performance detection, namely the Jaccard index between the real and detected perturbations, as a function of *α* and for different values of *β*.

## Discussion

How can we represent a temporal network, beyond a representation as a stream of interactions? This question can be answered differently depending on the system considered and on the goal of the representation.

For instance, recent proposals include static lossless representations of temporal networks, notably the supra-adjacency representation method [Valdano et al., 2015] and the event-graph [Kivelä et al., 2018], in which nodes and interactions are suitably mapped onto the nodes and links of static networks. These representations have shown to be useful for embedding and prediction tasks [Sato et al., 2019, Torricelli et al., 2020].

Temporal aggregation procedures, on the other hand, lose temporal information but have provided in-depth knowledge on the dynamics of social networks at various timescales [Aledavood et al., 2015, Saramäki et al., 2014, Fournet and Barrat, 2014, Saramäki et al., 2014]. Aggregated networks are also used for data-driven numerical simulations of dynamic processes of networks [Stehlé et al., 2011a], possibly with aggregation schemes adapted to the specific process under study [Holme, 2013].

Here, we consider an alternative type of representation: namely, a transformation of the temporal network stream into an evolving weighted network, which aims at providing a representation of the social system at any time and a description of its dynamics. Crucially, this transformation takes into account the interdependence of ties and the limited resources of any individual through the following ingredients: any interaction between two individuals reinforces their common tie and weakens the ties they have with other individuals not involved in the interaction.

While these ingredients can be translated in various ways into specific rules of evolution, here we have focused on a parsimonious two-parameter model rather than on more complex alternatives. We have applied this model to several data sets of interest, showing its ability to highlight changes in the dynamics of the networks and differences between data representing interactions in different contexts. Moreover, we have systematically tested its ability to detect a perturbation in the network at different timescales. Notably, our results show that this simple model yields a high detection performance even for small and short perturbations that cannot be detected by the dynamics of successive aggregated networks. Overall, our framework is able to detect perturbations in a broad range of conditions spanning different data sets and various timescales and perturbations. This point is particularly important as real-world variations in social relationships can occur on a broad range of timescales, from hours to days to months. For instance, despite decades of research, the timescale of the exchange of favors in primates (e.g., grooming in exchange for other commodities) is still very uncertain [Sánchez-Amaro and Amici, 2015]. Our framework does not require an a priori specification of the timescale of changes to be detected, but a scan of the parameters can help find the natural timescale(s) of the system under scrutiny. To investigate this point in more detail, further research will use a collection of temporal network models with tunable parameters and different levels of complexity and realism [Perra et al., 2012,Laurent et al., 2015]. Introducing perturbations of various types (e.g., changes in the community structure over time, changes in activity, etc), and of tunable intensity and duration, will allow us to systematically explore the detection capacities and limitations of the evolving weighted graph framework that we have introduced here.

An interesting property of our framework is that, starting from a stream of undirected interactions, it yields directed ties, because individuals do not invest in their mutual relationship in the same way: for instance, one individual may spend 80% of her time with another, while the other spends only 50% of her time with the first). The weights on each tie can therefore be more or less symmetric, and it would be interesting to investigate the significance of this (a)symmetry with respect to the social relationships under study. To this aim, one would need to compare the directed network obtained from our framework to other independent measures, such as friendship surveys in a human group or grooming behavior in non-human primates.

While we have limited our current study to a simple version of the model, several extensions could be of interest. In particular, directed interactions between individuals (such as phone or online messages) could be taken into account, with different impacts on the ties originating from the source of the interaction and on the ties originating from the interaction target. Moreover, one could take into account individual characteristics that are often important in relationships by introducing *α* and *β* coefficients that depend on individual characteristics such as age, sex, kinship or rank. This would be appropriate for instance when the costs and benefits of interactions differ between low ranking and high ranking individuals [Silk et al., 1999].

It is also worth mentioning the concepts of social contagion, consensus formation and social influence as potential application fields of our framework [Guilbeault et al., 2018,Rosenthal et al., 2015]. Social influence and contagion models are typically considered either on static aggregated networks or on temporal networks, each interaction conveying a potential event of social contagion. However, interactions with different individuals are in fact not equivalent, and our framework could provide a natural way to dynamically weigh these interactions: an interaction along a currently strong tie could weigh more than along a weak tie. This could provide a social contagion counterpart to the concept of epidemiologically optimal static networks to feed data-driven models of infectious diseases [Holme, 2013].

Finally, our focus here has been on social relationships of primates in particular, but our conceptual contribution lies in taking into account the interdependence of ties in evolving networks. Thus, our framework may well apply to other systems where such interdependence is relevant, possibly with changes in the rules of evolution. In particular, we have considered that an interaction between two nodes reinforces the tie between them at the expense of ties with other nodes, but in other contexts, the increase of a tie’s weight may in fact increase the importance of related ties. For instance, if a new flight route is created between two airports, passengers may take other flights to connect to other destinations, increasing the traffic on the corresponding routes [Barrat et al., 2004]. Taking these interactions into account might open up new perspectives to study the evolution of these types of infrastructure networks [Sugishita and Masuda, 2020].

## Materials and Methods

### Data Description and Aggregation

We used three datasets of time-stamped dyadic interactions between individuals corresponding to physical proximity events:

- A dataset of contacts between students in an urban public middle school in Utah (USA) measured by an infrastructure based on wireless ranging enabled nodes (WRENs) [Toth et al., 2015, Leecaster et al., 2016]. The data, available in reference [Leecaster et al., 2016], involve 679 students in grades 7 and 8 (typical age range from 12 to 14 years old). Participants were recorded over two consecutive days.
- A data set gathered by the SocioPatterns collaboration (http://www.sociopatterns.org/) using radio-frequency identification devices in an elementary school in France. These sensors record face-to-face contacts within a distance of about 1.5m. The data were aggregated with a temporal resolution of 20 seconds (for more details see [Cattuto et al., 2010]): two individuals were defined as being in contact during a 20s time window if their sensors exchanged at least one packet during that interval, and the contact event was considered to be over when the sensors no longer exchanged packets over a 20s interval. Contacts between 242 participants (232 elementary school children and 10 teachers) were recorded over two consecutive days [Stehlé et al., 2011b]. The data are publicly available at http://www.sociopatterns.org/datasets.
- Data of proximity contacts within a group of Guinea baboons (*Papio papio*), collected from June to November 2019 using an ad-hoc system of wearable devices. A subgroup of 13 baboons consisting only of juveniles and adults (all individuals were at least 6 years old) were equipped with leather collars fitted with the wearable proximity sensors developed by the SocioPatterns collaboration (see [Gelardi et al., 2020] for details).

### Similarity between networks

To compare the weighted evolving networks (or aggregated networks) observed at different times, we chose the global cosine similarity between the two vectors formed by the list of all the weights in each network (using a weight 0 if a link was not present).

A cosine similarity measure is generally defined between two vectors and is bounded between −1 and +1. It takes the value 1 if the vectors are proportional with a positive proportionality constant, a value of −1 if the proportionality constant is negative, and 0 if they are perpendicular. For positive weights, as in our case, it is bounded between 0 and 1.

In the case of two networks, *G*_1_ and *G*_2_, the global cosine similarity is precisely defined as:

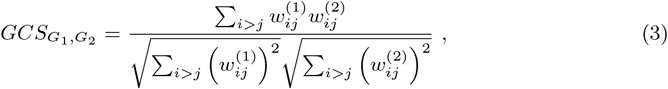

where the subscripts ^(1)^ and ^(2)^ denote the weights of the links in the networks *G*_1_ and *G*_2_, respectively.

### Clustering method

To obtain discrete system states by hierarchical clustering, we used the “fcluster” function of the scipy.hierarchy library from the SciPy module in Python. The function is applied directly on the *t_max_* × *t_max_* distance matrix *d*, obtained by transforming the cosine similarity matrix elements for each pair of timestamps (*t*, *t’*): *d*(*t*, *t’*) = 1 − *CS*(*t*, *t’*). To define the distance between clusters, we used the “average” method in the “linkage” function of the library. We set the number of clusters to *C* = 3, corresponding to the periods before, during and after the perturbation.

### Detection performance

Once we obtained the discrete states, we quantified the “quality” of the partition in order to decide which network representation (i.e. which value or set of values of the parameters) would be more appropriate to describe the system’s dynamics.

Our rationale was that the temporal network representation should allow us to detect changes in the social structure of the system under study, and the quality of the detection entails two aspects: it has to be detected (i) without delays and (ii) clearly, i.e., social changes have to be distinguished from the noise represented by “ordinary” variations in social activity. In particular, a perturbation is said to be well detected if one of the states found by the clustering algorithm includes all the timestamps of the perturbation and only those.

We first verified that one of the detected clusters could be associated with the perturbation in the data. To this end we determined that each cluster would correspond to a set of contiguous timestamps (thus forming an interval), with the smallest time equal to or larger than the initial timestamp of the perturbation, and largest time equal to or larger than the final timestamp of the perturbation. A first measure to evaluate the quality of the detection was then given by the “delay” between the actual and the detected perturbation (the number of timestamps between the actual starting time of the perturbation and the smallest timestamp of the second cluster detected; see Figure 2d). The second measure was given by the Jaccard index *J* between the set of time steps during which the actual perturbation takes place, 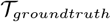, and the set of time steps of the state detected as a perturbation by the clustering procedure, 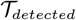:

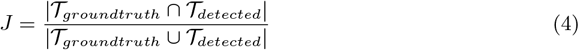

## Acknowledgments

Many thanks to Yousri Marzouki for planting the seed of the idea for this article and to Clément Sire for interesting discussions and the suggestion of studying case *β* ≠ *α* in the model. A.B. was supported by the ANR project DATAREDUX (ANR-19-CE46-0008) and JSPS KAKENHI Grant Number JP 20H04288.

## References

Ahmad, W., Porter, M. A., and Beguerisse-Díaz, M. (2018). Tie-decay temporal networks in continuous time and eigenvector-based centralities.

Aledavood, T., Lehmann, S., and Saramäki, J. (2015). Digital daily cycles of individuals. Frontiers in Physics, 3:73.

Barrat, A., Barthélemy, M., and Vespignani, A. (2004). Weighted evolving networks: Coupling topology and weight dynamics. Phys. Rev. Lett., 92:228701.

Brent, L. J., Lehmann, J., and Ramos-Fernández, G. (2011). Social network analysis in the study of nonhuman primates: A historical perspective. American Journal of Primatology, 73(8):720–730.

Burkart, J. M., Allon, O., Amici, F., Fichtel, C., Finkenwirth, C., Heschl, A., Huber, J., Isler, K., Kosonen, Z. K., Martins, E., Meulman, E. J., Richiger, R., Rueth, K., Spillmann, B., Wiesendanger, S., and van Schaik, C. P. (2014). The evolutionary origin of human hyper-cooperation. Nature Communications, 5(1):4747.

Cattuto, C., Van den Broeck, W., Barrat, A., Colizza, V., Pinton, J.-F., and Vespignani, A. (2010). Dynamics of person-to-person interactions from distributed RFID sensor networks. PLOS ONE, 5(7):1–9.

Cheney, D., Seyfarth, R., and Smuts, B. (1986). Social relationships and social cognition in nonhuman primates. Science, 234(4782):1361–1366.

Ciaperoni, M., Galimberti, E., Bonchi, F., Cattuto, C., Gullo, F., and Barrat, A. (2020). Relevance of temporal cores for epidemic spread in temporal networks. Scientific Reports, 10(1):12529.

Darst, R. K., Granell, C., Arenas, A., Gómez, S., Saramäki, J., and Fortunato, S. (2016). Detection of timescales in evolving complex systems. Scientific Reports, 6:39713.

Dunbar, R. (2020). Structure and function in human and primate social networks: Implications for diffusion, network stability and health. Proceedings of the Royal Society A, 476(2240):20200446.

Dunbar, R. and Shultz, S. (2017). Why are there so many explanations for primate brain evolution? Philosophical Transactions of the Royal Society B: Biological Sciences, 372(1727):20160244.

Dunbar, R. I. (1998). The social brain hypothesis. Evolutionary Anthropology: Issues, News, and Reviews: Issues, News, and Reviews, 6(5):178–190.

Dunbar, R. I., Korstjens, A. H., Lehmann, J., and Project, B. A. C. R. (2009). Time as an ecological constraint. Biological Reviews, 84(3):413–429.

Dunbar, R. I. and Shultz, S. (2007). Evolution in the social brain. science, 317(5843):1344–1347.

Eckmann, J. P., Moses, E., and Sergi, D. (2004). Entropy of dialogues creates coherent structures in e-mail traffic. PNAS, 101:14333–14337.

Elo, A. E. (1978). The rating of chessplayers, past and present. Arco Pub.

Fournet, J. and Barrat, A. (2014). Contact patterns among high school students. PLoS ONE, 9(9):e107878.

Galimberti, E., Barrat, A., Bonchi, F., Cattuto, C., and Gullo, F. (2018). Mining (maximal) span-cores from temporal networks. In Proceedings of the 27th ACM International Conference on Information and Knowledge Management, pages 107–116. ACM.

Gelardi, V., Fagot, J., Barrat, A., and Claidière, N. (2019). Detecting social (in)stability in primates from their temporal co-presence network. Animal Behaviour, 157:239–254.

Gelardi, V., Godard, J., Paleressompoulle, D., Claidière, N., and Barrat, A. (2020). Measuring social networks in primates: wearable sensors versus direct observations. Proceedings of the Royal Society A, 476:20190737.

Gonçalves, B., Perra, N., and Vespignani, A. (2011). Modeling users’ activity on twitter networks: Validation of dunbar’s number. PLOS ONE, 6(8):1–5.

Granovetter, M. S. (1973). The strength of weak ties. American Journal of Sociology, 78(6):1360–1380.

Granovetter, M. S. (1977). The strength of weak ties. In Social networks, pages 347–367. Elsevier.

Guilbeault, D., Becker, J., and Centola, D. (2018). Complex contagions: A decade in review. In Complex Spreading Phenomena in Social Systems, pages 3–25. Springer.

Heider, F. (1946). Attitudes and cognitive organization. The Journal of psychology, 21(1):107–112.

Hinde, R. A. (1976). Interactions, relationships and social structure. Man, 11(1):1–17.

Holme, P. (2013). Epidemiologically optimal static networks from temporal network data. PLoS Comput Biol, 9(7):e1003142.

Holme, P. (2015). Modern temporal network theory: a colloquium. The European Physical Journal B, 88(9):234.

Holme, P. and Saramäki, J. (2012). Temporal networks. Physics reports, 519(3):97–125.

Jin, E. M., Girvan, M., and Newman, M. (2001). Structure of growing social networks. Physical review. E, Statistical, nonlinear, and soft matter physics, 64 4 Pt 2:046132.

Karsai, M., Kivelä, M., Pan, R. K., Kaski, K., Kertész, J., Barabási, A.-L., and Saramäki, J. (2011). Small but slow world: How network topology and burstiness slow down spreading. Physical Review E, 83(2).

Karsai, M., Perra, N., and Vespignani, A. (2014). Time varying networks and the weakness of strong ties. Scientific Reports, 4(1):4001.

Kivelä, M., Cambe, J., Saramäki, J., and Karsai, M. (2018). Mapping temporal-network percolation to weighted, static event graphs. Scientific Reports, 8(1):12357.

Kivelä, M. and Porter, M. A. (2015). Estimating interevent time distributions from finite observation periods in communication networks. Physical Review E, 92(5):052813.

Kobayashi, T., Takaguchi, T., and Barrat, A. (2019). The structured backbone of temporal social ties. Nature Communications, 10(1):220.

Kossinets, G. and Watts, D. J. (2006). Empirical analysis of an evolving social network. Science, 311:88–90.

Kovanen, L., Karsai, M., Kaski, K., Kertész, J., and Saramäki, J. (2011). Temporal motifs in time-dependent networks. Journal of Statistical Mechanics: Theory and Experiment, 2011(11):P11005.

Kovanen, L., Kaski, K., Kertész, J., and Saramäki, J. (2013). Temporal motifs reveal homophily, gender-specific patterns, and group talk in call sequences. Proceedings of the National Academy of Sciences, 110(45):18070–18075.

Krings, G., Karsai, M., Bernhardsson, S., Blondel, V. D., and Saramäki, J. (2012). Effects of time window size and placement on the structure of an aggregated communication network. EPJ Data Science, 1(1):4.

Kwak, S., Joo, W.-t., Youm, Y., and Chey, J. (2018). Social brain volume is associated with in-degree social network size among older adults. Proceedings of the Royal Society B: Biological Sciences, 285(1871):20172708.

Laurent, G., Saramäki, J., and Karsai, M. (2015). From calls to communities: a model for time-varying social networks. The European Physical Journal B, 88(11):301.

Leecaster, M., Toth, D. J. A., Pettey, W. B. P., Rainey, J. J., Gao, H., Uzicanin, A., and Samore, M. (2016). Estimates of social contact in a middle school based on self-report and wireless sensor data. PLOS ONE, 11(4):1–21.

Lewis, P. A., Rezaie, R., Brown, R., Roberts, N., and Dunbar, R. I. (2011). Ventromedial prefrontal volume predicts understanding of others and social network size. Neuroimage, 57(4):1624–1629.

Masuda, N. and Holme, P. (2019). Detecting sequences of system states in temporal networks. Scientific reports, 9(1):1–11.

Meguerditchian, A., Marie, D., Margiotoudi, K., Roth, M., Nazarian, B., Anton, J.-L., and Claidière, N. (2021). Baboons (papio anubis) living in larger social groups have bigger brains. Evolution and Human Behavior, 42:1–30.

Miritello, G., Lara, R., Cebrian, M., and Moro, E. (2013a). Limited communication capacity unveils strategies for human interaction. Scientific reports, 3(1):1–7.

Miritello, G., Lara, R., and Moro, E. (2013b). Time allocation in social networks: correlation between social structure and human communication dynamics. In Temporal networks, pages 175–190. Springer.

Mitani, J. C. (2009). Male chimpanzees form enduring and equitable social bonds. Animal Behaviour, 77(3):633–640.

Moreno, J. L. (1934). Who Shall Survive? Foundations of Sociometry, Group Psychotherapy, and Sociodram. Beacon House, NY.

Navarro, H., Miritello, G., Canales, A., and Moro, E. (2017). Temporal patterns behind the strength of persistent ties. EPJ Data Science, 6(1):31.

Noonan, M., Mars, R., Sallet, J., Dunbar, R., and Fellows, L. (2018). The structural and functional brain networks that support human social networks. Behavioural brain research, 355:12–23.

Onnela, J.-P., Saramäki, J., Hyvönen, J., Szabó, G., Lazer, D., Kaski, K., Kertész, J., and Barabási, A.-L. (2007). Structure and tie strengths in mobile communication networks. Proceedings of the National Academy of Sciences, 104(18):7332–7336.

Palla, G., Barabási, A.-L., and Vicsek, T. (2007). Community dynamics in social networks. Fluctuation and Noise Letters, 07(03):L273–L287.

Perra, N., Gonçalves, B., Pastor-Satorras, R., and Vespignani, A. (2012). Activity driven modeling of time varying networks. Scientific Reports, 2(1):469.

Psorakis, I., Roberts, S. J., Rezek, I., and Sheldon, B. C. (2012). Inferring social network structure in ecological systems from spatio-temporal data streams. Journal of the Royal Society Interface, 9(76):3055–3066.

Rosenthal, S. B., Twomey, C. R., Hartnett, A. T., Wu, H. S., and Couzin, I. D. (2015). Revealing the hidden networks of interaction in mobile animal groups allows prediction of complex behavioral contagion. Proceedings of the National Academy of Sciences, 112(15):4690–4695.

Salathé, M., Kazandjieva, M., Lee, J. W., Levis, P., Feldman, M. W., and Jones, J. H. (2010). A high-resolution human contact network for infectious disease transmission. Proceedings of the National Academy of Sciences, 107(51):22020–22025.

Saramäki, J., Leicht, E. A., López, E., Roberts, S. G. B., Reed-Tsochas, F., and Dunbar, R. I. M. (2014). Persistence of social signatures in human communication. Proceedings of the National Academy of Sciences, 111(3):942–947.

Saramäki, J. and Moro, E. (2015). From seconds to months: an overview of multi-scale dynamics of mobile telephone calls. The European Physical Journal B, 88(6):164.

Sato, K., Oka, M., Barrat, A., and Cattuto, C. (2019). Dyane: Dynamics-aware node embedding for temporal networks.

Silk, J., Cheney, D., and Seyfarth, R. (1999). The structure of social relationships among female savanna baboons in moremi reserve, botswana. Behaviour, 136:679–703.

Silk, J. B., Beehner, J. C., Bergman, T. J., Crockford, C., Engh, A. L., Moscovice, L. R., Wittig, R. M., Seyfarth, R. M., and Cheney, D. L. (2010). Female chacma baboons form strong, equitable, and enduring social bonds. Behavioral Ecology and Sociobiology, 64(11):1733–1747.

Stehlé, J., Voirin, N., Barrat, A., Cattuto, C., Colizza, V., Isella, L., Régis, C., Pinton, J.-F., Khanafer, N., Van den Broeck, W., et al. (2011a). Simulation of an seir infectious disease model on the dynamic contact network of conference attendees. BMC medicine, 9(1):87.

Stehlé, J., Voirin, N., Barrat, A., Cattuto, C., Isella, L., Pinton, J.-F., Quaggiotto, M., Van den Broeck, W., Régis, C., Lina, B., et al. (2011b). High-resolution measurements of face-to-face contact patterns in a primary school. PloS one, 6(8):e23176.

Stopczynski, A., Sekara, V., Sapiezynski, P., Cuttone, A., Madsen, M. M., Larsen, J. E., and Lehmann, S. (2014). Measuring large-scale social networks with high resolution. PloS one, 9(4):e95978.

Sugishita, K. and Masuda, N. (2020). Recurrence in the evolution of air transport networks.

Sulo, R., Berger-Wolf, T., and Grossman, R. (2010). Meaningful selection of temporal resolution for dynamic networks. In Proceedings of the Eighth Workshop on Mining and Learning with Graphs, pages 127–136.

Szell, M., Lambiotte, R., and Thurner, S. (2010). Multirelational organization of large-scale social networks in an online world. Proceedings of the National Academy of Sciences, 107:13636.

Sánchez-Amaro, A. and Amici, F. (2015). Are primates out of the market? Animal Behaviour, 110:51–60.

Taebi, A., Kiesow, H., Vogeley, K., Schilbach, L., Bernhardt, B. C., and Bzdok, D. (2020). Population variability in social brain morphology for social support, household size, and friendship satisfaction. Social Cognitive and Affective Neuroscience, 15:635–647.

Tomasello, M., Carpenter, M., Call, J., Behne, T., and Moll, H. (2005). Understanding and sharing intentions: the origins of cultural cognition. Behavioral and Brain Sciences, 28(5):675–91; discussion 691–735.

Torricelli, M., Karsai, M., and Gauvin, L. (2020). weg2vec: Event embedding for temporal networks. Scientific Reports, 10(1):7164.

Toth, D. J., Leecaster, M., Pettey, W. B., Gundlapalli, A. V., Gao, H., Rainey, J. J., Uzicanin, A., and Samore, M. H. (2015). The role of heterogeneity in contact timing and duration in network models of influenza spread in schools. Journal of The Royal Society Interface, 12(108):20150279.

Ureña-Carrion, J., Saramäki, J., and Kivelä, M. (2020). Estimating tie strength in social networks using temporal communication data. EPJ Data Science, 9(1):37.

Valdano, E., Ferreri, L., Poletto, C., and Colizza, V. (2015). Analytical computation of the epidemic threshold on temporal networks. Phys. Rev. X, 5:021005.

Wasserman, S. and Faust, K. (1994). Social Network Analysis: Methods and applications. Cambridge University Press, Cambridge.

Zuo, X. and Porter, M. A. (2019). Models of continuous-time networks with tie decay, diffusion, and convection.

