## Supplemental figures for "From temporal network data to the dynamics of social relationships"

### Supporting Information

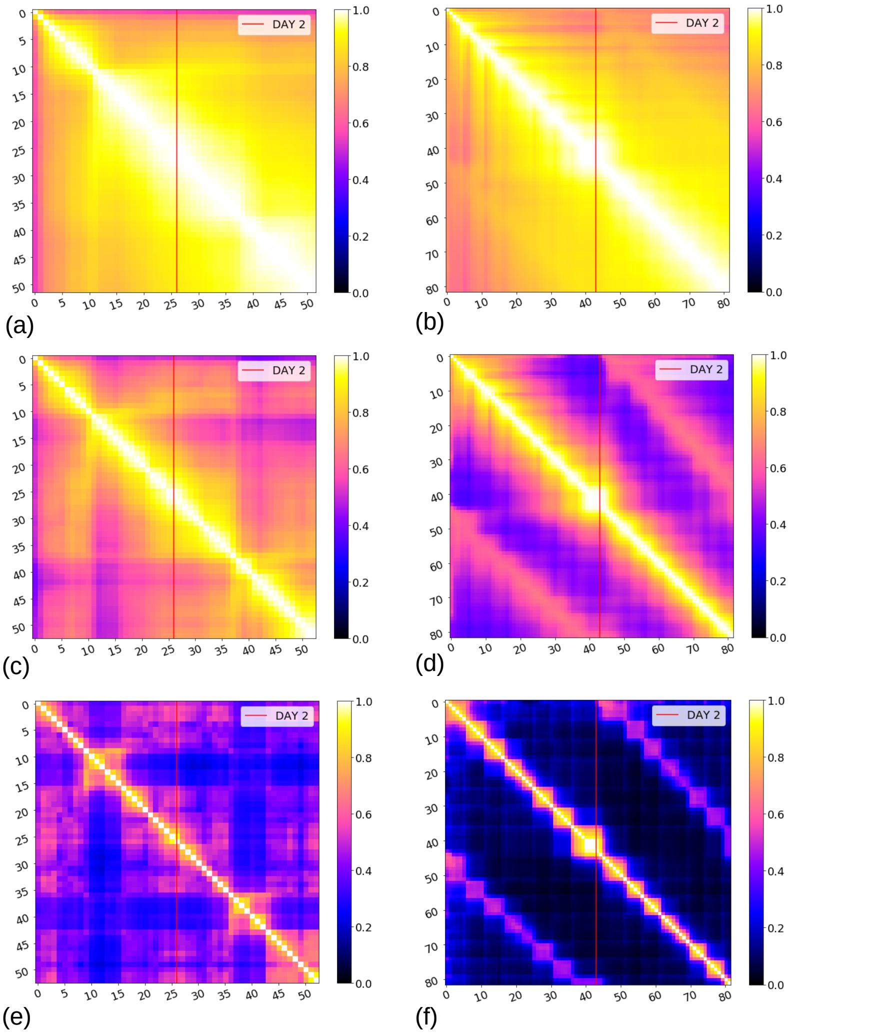


**Figure S1. Cosine Similarity matrices for the French elementary school and Utah middle school, using different values of $\alpha$.** The first column (a),(c),(e) refers to the French elementary school data, sampling the network every 20 minutes; the second column refers to the Utah middle school data, sampling the network every 10 minutes. The global cosine similarity is calculated between every pair of observed networks. The vertical red lines indicate the time between the first and the second day of data collection (the data of the two days are concatenated). The different rows refer to three different values of the parameter $\alpha$, namely $\alpha = 0.001$ (a) (b), $\alpha = 0.01$ (c) (d) and $\alpha = 0.1$ (e)(f). For $\alpha = 0.1$ the different structures of the schools schedules emerge more clearly. We observe for the French school that the structure of the networks during the two lunches are different, and that the structure during the classes remain similar in the two days. For the Utah middle school, we observe some similarity between the corresponding class periods in the two different days, indicating that the seating arrangements in each class are probably similar in different days.

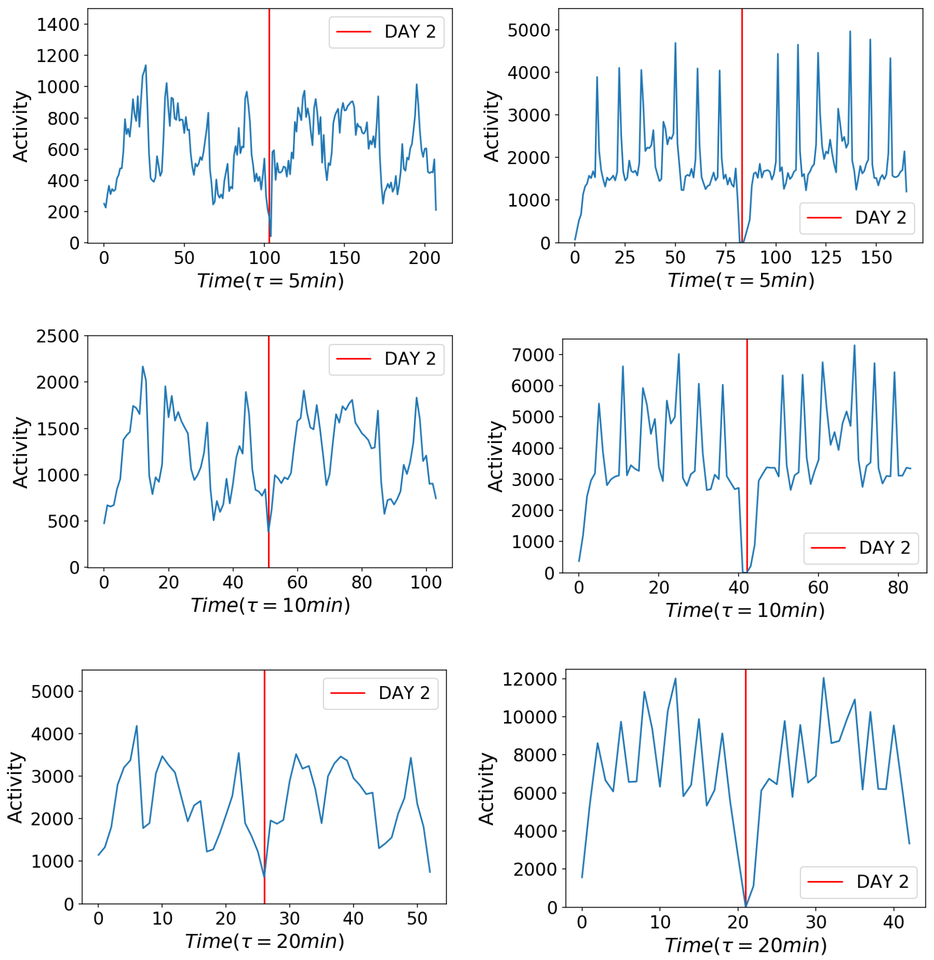


**Figure S2. Event rate (number of temporal edges per timestamp) vs. time** for the French elementary school data (left column) and the Utah middle school data (right column). Each row corresponds to a different time resolution (respectively 5, 10 and 20 minutes). While the activity timelines can highlight some timestamps of relevance, they do not shed information on the evolution of the structure of the system (i.e., whether the networks at different times have similar or different structures).

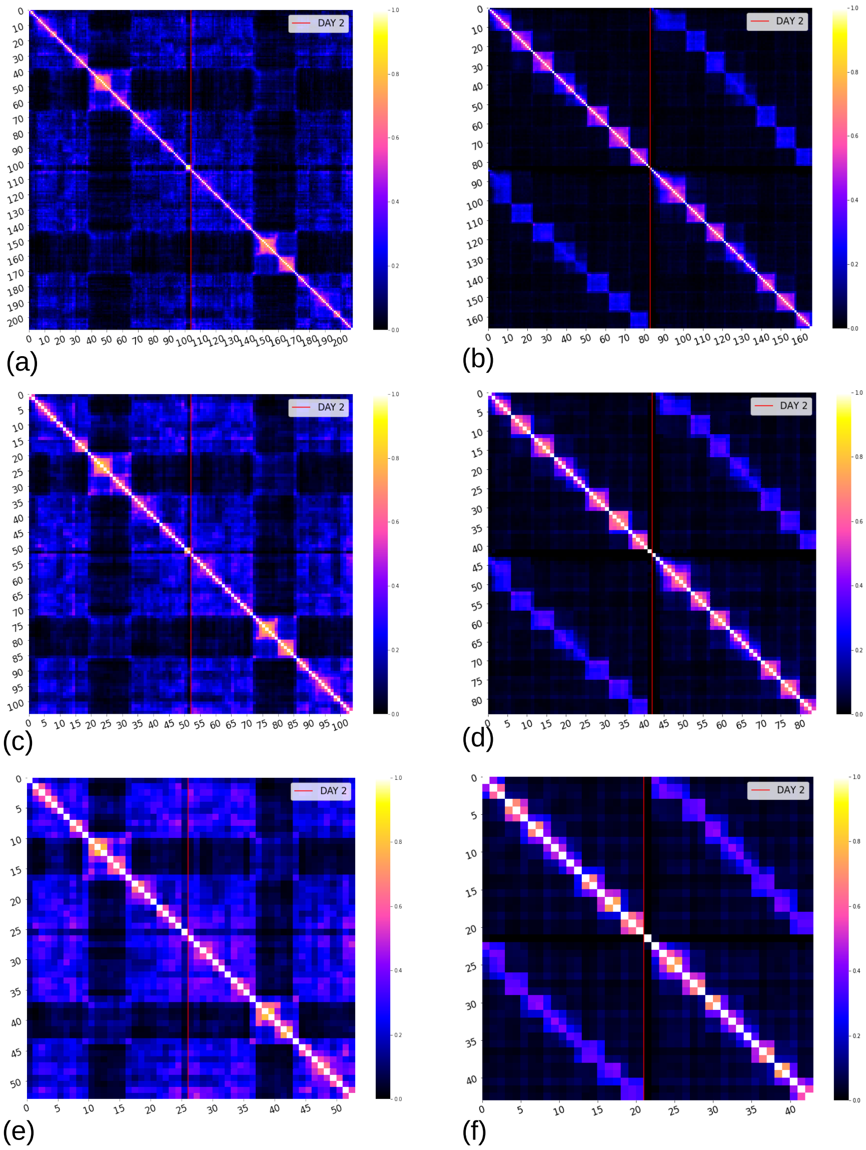


**Figure S3. Cosine Similarity matrices for the French elementary school and Utah middle school, aggregating the network over different time window lengths.** The first column (a),(c),(e) refers to the French elementary school data ; the second column refers to the Utah middle school data. The different rows refer to three different values of the aggregation time window length: respectively 5, 10 and 20 minutes.

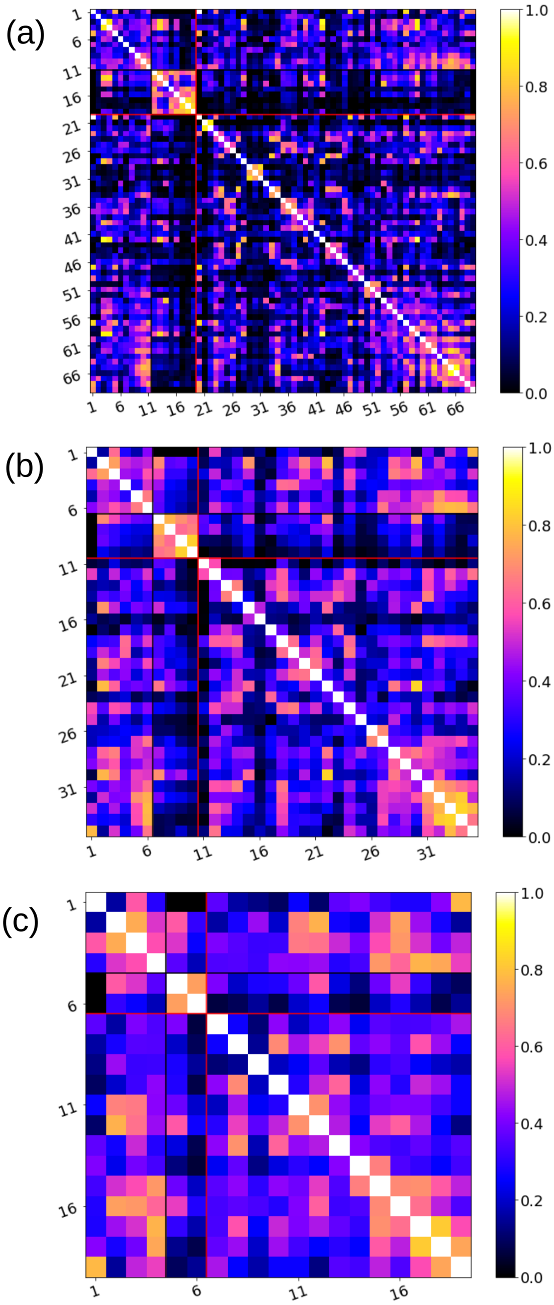


**Figure S4. Cosine Similarity matrices for the baboon data, aggregating the network over different time scales.** The three panels refer to the aggregated networks computed using the same data as in Fig. 3 for three different values of the aggregation time window, i.e. 15 minutes (a), 30 minutes (b) and 1 hour (c).

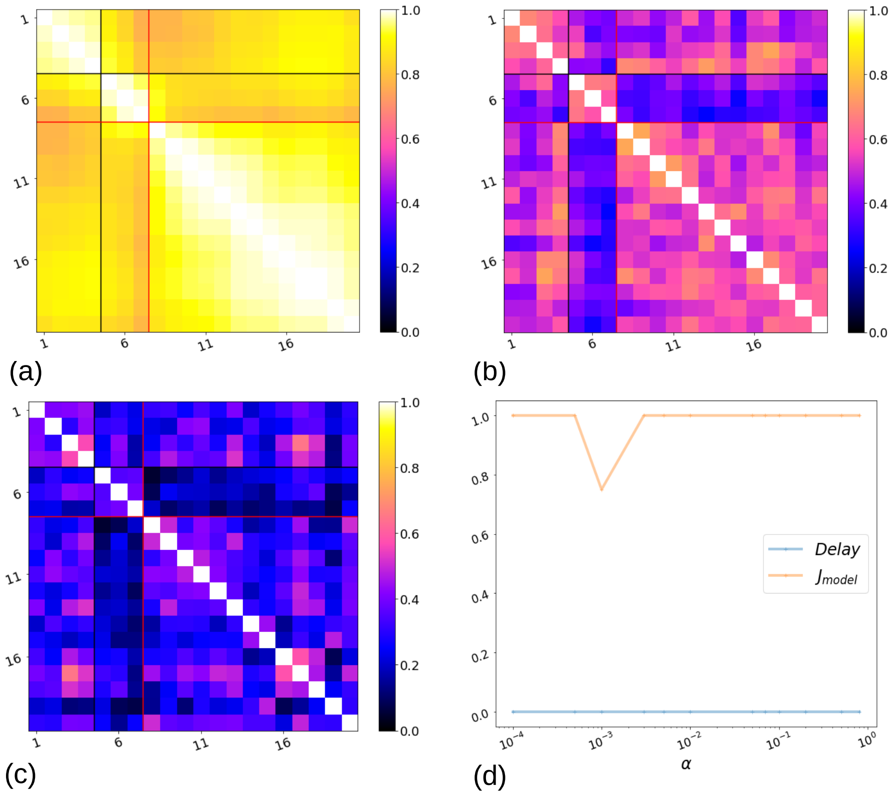


**Figure S5.** Detection of a simulated perturbation in a temporal network data set. Here we consider 20 days of proximity data collected in a group of 13 baboons (see Material and Methods). The data, whose temporal resolution is 20 seconds, is artificially perturbed by exchanging the identity of two nodes for 3 days. The resulting perturbed temporal network is transformed into a weighted evolving network as described in the text, and this network is observed on a daily basis. Panels (a), (b), (c) represent the resulting cosine similarity matrices for values of $\alpha = \beta = 0.001$, 0.1, 0.5, respectively. The black and red lines correspond to the (known) start and end times of the perturbation. Panel (d) shows the detection performance (see Fig. 2), computed from the hierarchical clustering analysis applied to the distance matrices, with the number of clusters fixed to $C = 3$. The blue line represents the relative delay in the detection of the perturbation, i.e. the difference between the known beginning of the perturbation (black line) and the detection of a new network state, divided by the total length of the perturbation. The orange line indicates the Jaccard index between the known perturbation timestamps and the perturbation detected by the clustering algorithm using different values of the parameter $\alpha$. The detection performance relative to the aggregated network is missing because no cluster detected by the algorithm could correspond to the simulated perturbation.

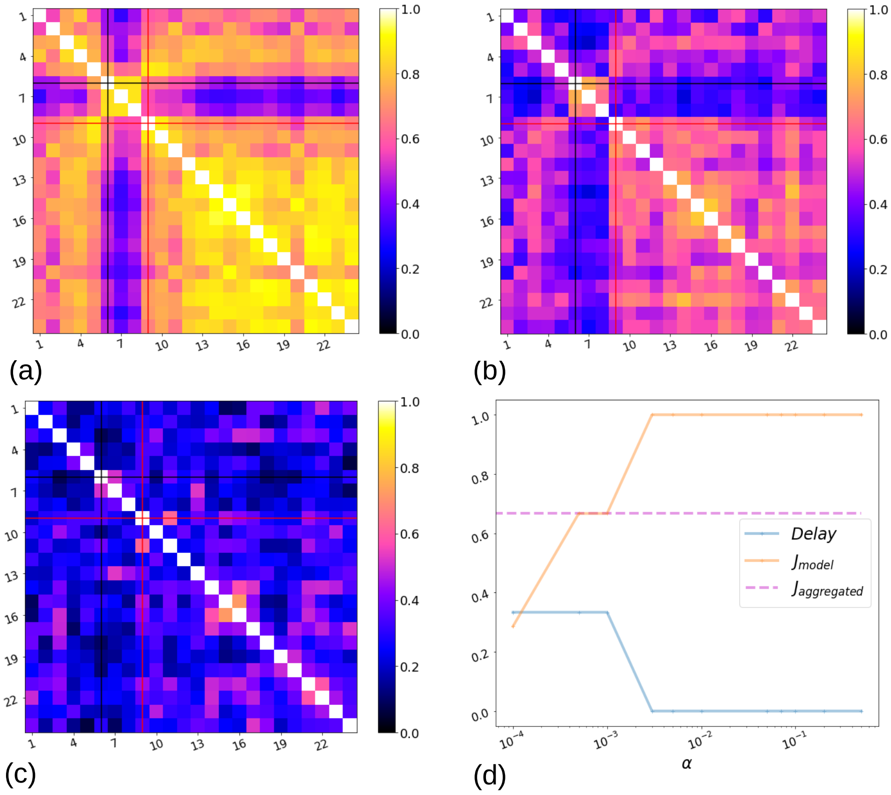


**Figure S6.** Detection of a simulated perturbation in a temporal network data set. Here we consider the proximity data collected in a group of 13 baboons over several months (see Material and Methods). The data, whose temporal resolution is 20 seconds, is artificially perturbed by exchanging the identity of two nodes during 15 days, affecting weeks 6 to 8. The resulting perturbed temporal network is transformed into a weighted evolving network as described in the text, and this network is here observed every 7 days. Panels (a), (b), (c) represent the resulting cosine similarity matrices for values of $\alpha = \beta = 0.001$, 0.1, 0.5, respectively. The black and red lines correspond to the (known) start and end times of the perturbation. Panel (d) shows the detection performance (see Fig. 2), computed from the hierarchical clustering analysis applied to the distance matrices, with the number of clusters fixed to $C = 3$. The blue line represents the relative delay in the detection of the perturbation, i.e. the difference between the known beginning of the perturbation (black line) and the detection of a new network state, divided by the total length of the perturbation. The orange line indicates the Jaccard index between the known perturbation timestamps and the perturbation detected by the clustering algorithm using different values of the parameter $\alpha$; the dotted magenta line indicates the Jaccard index using networks aggregated over successive 7 day time windows.

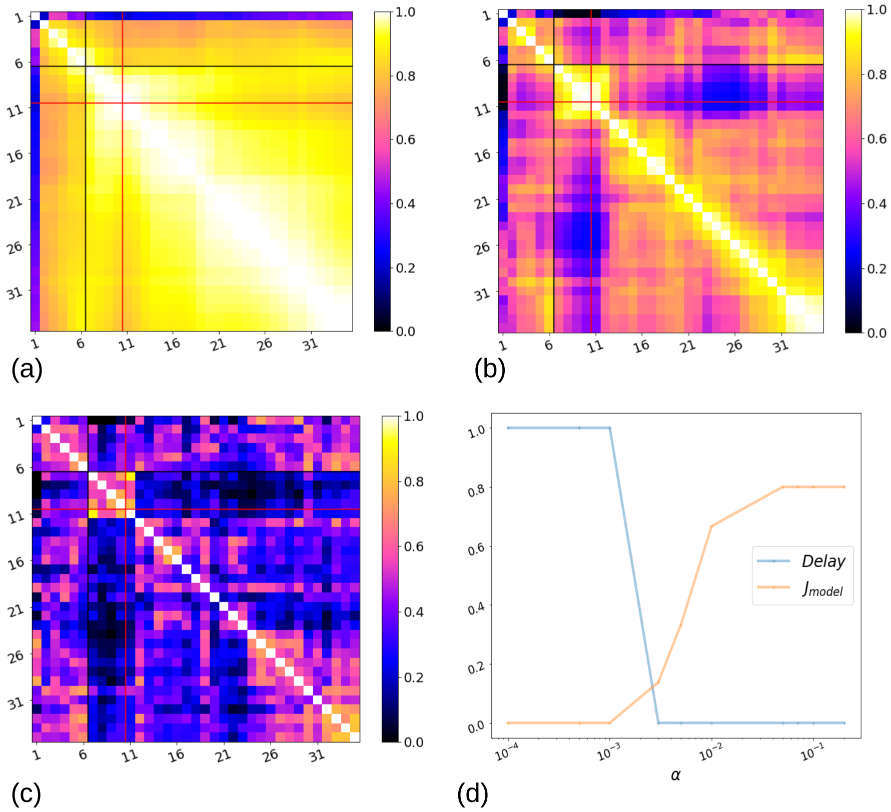


**Figure S7. Detection of simulated perturbations in the baboon' data set, using the weighted evolving network with different values of the parameters $\alpha$ and $\beta$.** Panels (a), (b), (c) represent the cosine similarity matrices for $\beta = 5\alpha$ and values of $\alpha = 0.001, 0.01$ and $0.1$, respectively. The networks and the detection performance were computed using the same procedure as in Fig. 3.
